# Splenic MARCO^+^ marginal zone macrophages regulate rapid production of MCP-1 and KC but are dispensable for alloantibody generation in response to stored RBCs in a murine model

**DOI:** 10.1101/2025.02.11.637644

**Authors:** Abhinav Arneja, Juan E. Salazar, Jelena Medved, Sarah J. Ratcliffe, Mark E. Smolkin, Manjula Santhanakrishnan, Sean R. Stowell, Krystalyn E. Hudson, James C. Zimring, Jeanne E. Hendrickson, Chance John Luckey

## Abstract

**BACKGROUND:** Alloimmunization to transfused red blood cells (RBCs) remains a significant clinical problem. However, the cells that initiate immune responses to transfused RBCs remain incompletely characterized.

Recently published work has identified splenic marginal zone B (MZB) cells as being critically required for the production of anti-RBC alloantibodies in response to RBCs. In infectious models, MZB cell activation has been shown to depend on a unique population of marginal zone macrophages (MZMs). We hypothesized that MZMs would capture stored RBCs and present them to MZBs, and ultimately MZMs would be required for generation of anti-RBC alloantibodies in response to stored RBC transfusion.

**STUDY DESIGN AND METHODS:** Stored GFP^+^ murine RBCs were utilized to determine the splenic localization and erythrophagocytosis by splenic macrophage populations. To determine the functional impact of MZMs, we compared LXRα-KO mice, which have been reported to lack MZMs, with wild type mice. Both innate and adaptive immune responses to stored HOD allogenic RBC transfusion were measured in LXRα-KO and wild type mice.

**RESULTS:** RBC storage leads to a significant increase in the phagocytosis of transfused RBCs by splenic MZMs. LXRα-KO mice demonstrated a lack of MZMs and had significantly decreased rapid phase production of cytokines MCP-1 and KC, but similar levels of IL-6. Surprisingly, anti-RBC alloantibody levels were unaffected by the absence of splenic MZMs.

**CONCLUSIONS:** Splenic MZMs are involved in the innate response to transfused stored HOD RBCs, contributing to both MCP-1 and KC cytokine production. However, MZMs are dispensable for anti-RBC alloantibody production.

## INTRODUCTION

Alloimmunization to non-ABO blood group antigens on transfused red blood cells (RBCs) remains a significant problem in transfusion medicine. Re-exposure to the same RBC antigens in the presence of previously generated alloantibodies can lead to hemolytic transfusion reactions with the potential for causing significant morbidity and occasional mortality in alloimmunized patients^1^. Presence of alloantibodies can make location, testing, and provision of life-saving transfusion therapies difficult or, in rare cases, impossible. Despite the clinical significance of RBC alloimmunization, our understanding of the mechanisms by which allogenic RBC transfusion drives innate cell activation remains limited^1,2^.

Mice engineered to express alloantigens on the surface of their RBCs have provided novel experimentally tractable model systems which allow for investigation of the cellular and molecular mechanisms of RBC alloimmunization. Since all allogenic human RBCs undergo some period of refrigerator storage prior to transfusion, we and others have previously investigated the role that refrigerated storage has on the resulting immune response to transfused blood. Transfusion of mouse blood that has been stored at the extremes of the FDA-allowable storage duration drives the rapid production of multiple inflammatory cytokines^3,4^. Transfusion of stored RBCs correlates with significantly reduced recovery post-transfusion compared to fresh RBCs, suggesting increased interactions with cellular compartments that result in their removal from circulation.^3^ In a study by Wojczyk et al., physiological depletion of phagocytes significantly reduced the rapid cytokine production in response to transfused stored RBCs in mice, and the use of reporter mice showed splenic tissue resident macrophages as the cells responsible for the production of inflammatory cytokines in response to stored RBC transfusion^5^. Collectively, these results suggest an important role for various splenic phagocyte populations as the initiators of the innate immune response generated by stored RBC transfusion. However, the specific splenic phagocyte populations that are responsible for rapid cytokine production in response to stored RBCs remains unclear.

Transfusion of stored RBCs has also been shown to increase anti-RBC alloantibody production in at least one mouse model of alloimmunization. Transfusion of RBCs expressing the triple fusion HOD antigen (<u>h</u>en egg lysozyme, <u>o</u>valbumin, and the human <u>D</u>uffy^b^ antigen) leads to enhancement of anti-RBC IgG alloantibodies relative to those generated in response to fresh RBCs^4,6^. Previous work has shown the spleen to be a critically important organ for RBC alloimmunization in both humans and mice, suggesting that splenic cell populations are required for RBC alloimmunization^7,8^. Within the spleen, splenic marginal zone B (MZB) cells have been shown to be critically required for the production of anti-RBC alloantibodies in response to stored HOD RBCs^9^. However, the mechanisms by which MZB support anti-RBC alloantibody formation in the HOD mouse model is unknown. While it’s certainly possible that MZB themselves directly produce anti-RBC alloantibodies, the mechanisms by which they are activated by stored blood transfusion remain to be discovered.

MZBs localize to the outer margins of the splenic sinus in mice where they come in contact with blood before it filters through to the red pulp^10,11^. In infectious models, MZB cell activation has been shown to depend on their interactions with a unique population of macrophages that co-localize with MZBs in the outer margin of the splenic sinus^12,13^. These marginal zone macrophages (MZMs) express the scavenger receptor MARCO^+^ and the C-type lectin SignR1^+^, and have been shown to play an important role in the capture of blood borne antigens, facilitating recognition of these antigens by MZBs. MZM macrophages are also known to trap, phagocytose, and initiate innate immune reactions to intravenously injected apoptotic cells^11,14,15^. Refrigerator storage of RBCs leads to myriad biophysical and biochemical changes^16–18^. Some of these changes include properties shared by apoptotic nucleated cells such as elevated intracellular Ca^2+^ and increased surface phosphatidlyserine (PS) levels^19,20^. A study by Larsson et al. showed that ionomycin treated murine RBCs, which had increased intracellular Ca^2+^ and surface PS levels, were trapped in the splenic marginal zone and showed significant co-localization with splenic MZMs compared to untreated RBCs^20^. Based on these results, we hypothesized that MZMs in the splenic marginal zone trap and phagocytose transfused refrigerator-stored RBCs. We further hypothesized that MZMs are involved in the rapid generation of inflammatory cytokines that have been shown to be upregulated in response to stored blood transfusion. Finally, we hypothesized that MZMs play a key role in the activation of MZBs in response to stored blood transfusion, driving MZB activation and ultimately leading to anti-RBC alloantibody production.

## MATERIALS AND METHODS

### Mice

C57BL/6J, FVB/NJ, B6.GFP (C57BL/6-Tg(UBC-GFP)30Scha/J), and LXRα-KO (B6;129S6-Nr1h3^tm1Djm^/J) mice were purchased from Jackson Laboratory (Bar Harbor, ME). HOD mice, on an FVB background, were generated as previously described^21^. All mice were house in pathogen free facilities at the University of Virginia and mouse protocols were approved by the Institutional Animal Care and Use Committees of Harvard Medical School and University of Virginia, Charlottesville.

### Murine blood collection, storage, and transfusion

Collection and processing of FVB.HOD and B6.GFP blood was performed as described previously^22^ and detailed in supplementary methods.

### Phagocyte depletion through clodronate liposome treatment

WT C57BL/6 mice were treated with clodronate loaded liposomes (clorodronateliposomes.com) through intravenous injections 48-72h prior to transfusion with stored HOD RBCs. Each animal received an equivalent of 2mg clodronate through the administration of clodronate liposomes.

### Immunofluorescence microscopy

Immunofluorescence staining and imaging of splenic macrophage populations and splenic localization of transfused GFP RBCs was performed as previously published^9,23^ and detailed in supplementary methods.

### Flow cytometry

Erythrocyte phagocytosis assay, MZ B cell staining, and MZM staining were performed as previously published^9,24^ and detailed in supplementary methods.

### Serum cytokine and antibody measurements

Sera from recipient mice were collected via submandibular bleeding 90 minutes post-transfusion for cytokine measurements; and at 7 and 14 days for antibody measurements as previously described^22^. Cytokine measurements, anti-HOD antibody flow crossmatch measurements, and anti-HEL ELISA measurements are detailed in supplementary methods.

### T independent antigen immunization and serum immunoglobulin measurements

WT C57BL/6 and LXRα-KO were immunized with 100μg of hapten 2,4,6-Trinitrophenol conjugated Ficoll (TNP-Ficoll, Biosearch Technologies) in PBS as previously published^23^ and detailed in supplemental methods.

### Statistical Analyses

All data were compared using Mann-Whitney U test with P values less than 0.05 considered as significant. For comparisons of more than one group, one-way Kruskal-Wallis analysis of variance test was initially performed followed by Mann-Whitney U test for post-hoc comparisons. All statistical tests were performed using GraphPad Prism software. To determine the impacts of cytokine levels across experiments, we further analyzed all the replicates together using generalized estimating equation (GEE) statistical analysis to account for variance introduced from each independent study (see supplementary methods).

## RESULTS

### Phagocytes are required for innate cell cytokine production and subsequent adaptive alloantibody generation in response to transfusion of stored RBCs

To measure antigen specific anti-RBC antibody responses, we used the well characterized HOD mouse model of RBC alloimmunization, in which mice express a fusion protein composed of hen egg lysozyme, ovalbumin, and the human Duffy blood group antigen on their RBCs^21^. We and others have previously shown that transfusion with stored RBCs leads to rapid production of multiple pro-inflammatory cytokines and enhanced anti-HOD alloantibody production^3,6,22^. To investigate the role of phagocytes in alloantibody production in response to transfused stored HOD RBCs, WT C57BL/6 mice were treated with clodronate liposomes (CLs) prior to transfusion with 12-day stored HOD RBCs. Clodronate loaded liposomes are preferentially phagocytosed by macrophages, resulting in phagocyte apoptosis^25^. Recipient mice treated with CLs showed an increased recovery post-transfusion and a significant reduction in serum levels of IL-6 and MCP-1 90 minutes post-transfusion with stored HOD RBCs (Figure 1A-B, supplemental fig. 1). This demonstrates that the initial cytokine burst observed in response to stored RBC transfusion depends on CL sensitive cells, likely macrophages.

**Figure 1.**
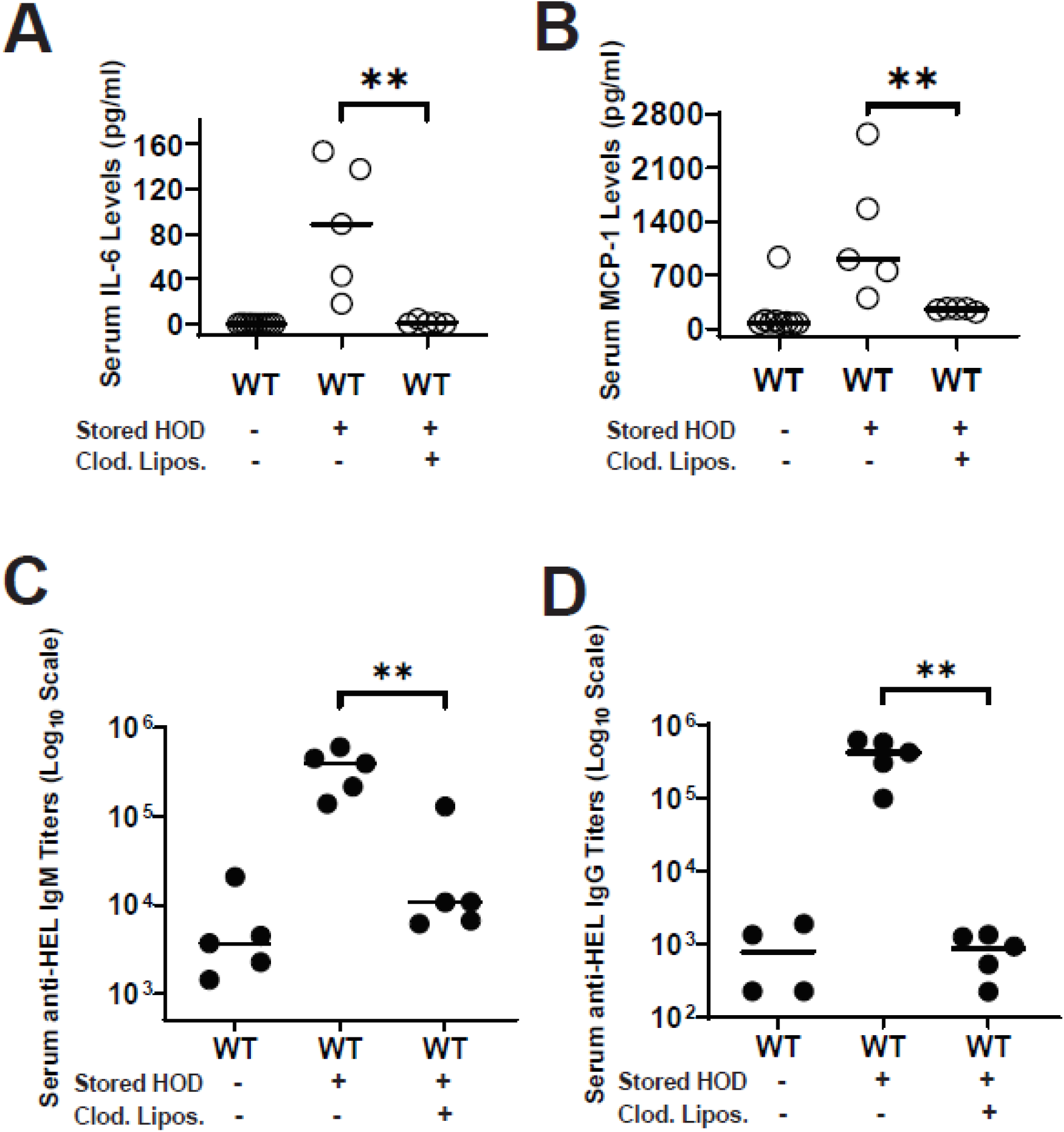
Phagocytes are required for rapid inflammatory cytokine production and alloantibody generation in response to stored HOD RBC transfusion. WT C57BL/6 mice, 8-12 weeks of age, were treated with clodronate liposomes to deplete phagocytes, and then transfused with 12-day stored HOD RBCs. Sera were collected at multiple times post-transfusion to measure serum levels of inflammatory cytokines and anti-HOD alloantibody levels. Serum levels of (A) IL-6 and (B) MCP-1 90min post-transfusion were measured and quantified through cytokine specific ELISAs. (C) Serum anti-HOD IgM titers 7 days post-transfusion measured through anti-HEL ELISAs, depicted on a log10 scale. (D) Serum anti-HOD IgG titers 14 days post-transfusion measured through anti-HEL ELISAs, depicted on a log10 scale. Data shown are representative of 3 independent experiments. Statistical significance was determined through non-parametric the Mann-Whitney U test. *p < 0.05, **p < 0.01, ***p < 0.001and ns p > 0.05.

Additionally, mice treated with CLs prior to transfusion had a significant reduction in serum anti-HEL IgM titers (Figure 1C) as well serum anti-HEL IgG titers (Figure 1D). Importantly, the impact of CL on IgM suggests that the initial B cell activation in response to RBC transfusion also depends on a CL sensitive cell type. These results suggest that both rapid cytokine production and alloantibody generation in response to transfusion of stored HOD RBCs appears to require liposome phagocytic activity, most likely macrophages.

### Stored RBCs localize to and are phagocytosed by marginal zone macrophages (MZM)

To determine if stored RBCs interact with MZMs, we first determined the splenic localization of transfused RBCs. Spleens from recipient mice transfused with stored GFP^+^ RBCs were collected at 60min post-transfusion, and imaged for GFP^+^ RBCs and MARCO^+^ MZMs. We found that stored RBCs clearly co-localized with splenic MZMs (Figure 2). It should be noted that we also observed significant GFP signal co-localizing with the other major splenic macrophage populations (Supplemental Figure 2). Next, we asked whether the stored RBCs were phagocytosed by MZMs. Spleens from recipient mice transfused with stored GFP^+^ RBCs were processed into single-cell suspension, then stained for MZMs using SIGNR1. Phagocytosis of GFP^+^ RBCs was determined through quantification of GFP^+^ macrophages via flow cytometry. We found that transfusion with stored GFP^+^ RBCs led to a clear subset of SIGNR1^+^ MZMs that were GFP positive (Figure 3, supplemental figure 3). These results show that refrigerator stored RBCs localize to and are phagocytosed by splenic MZMs.

**Figure 2.**
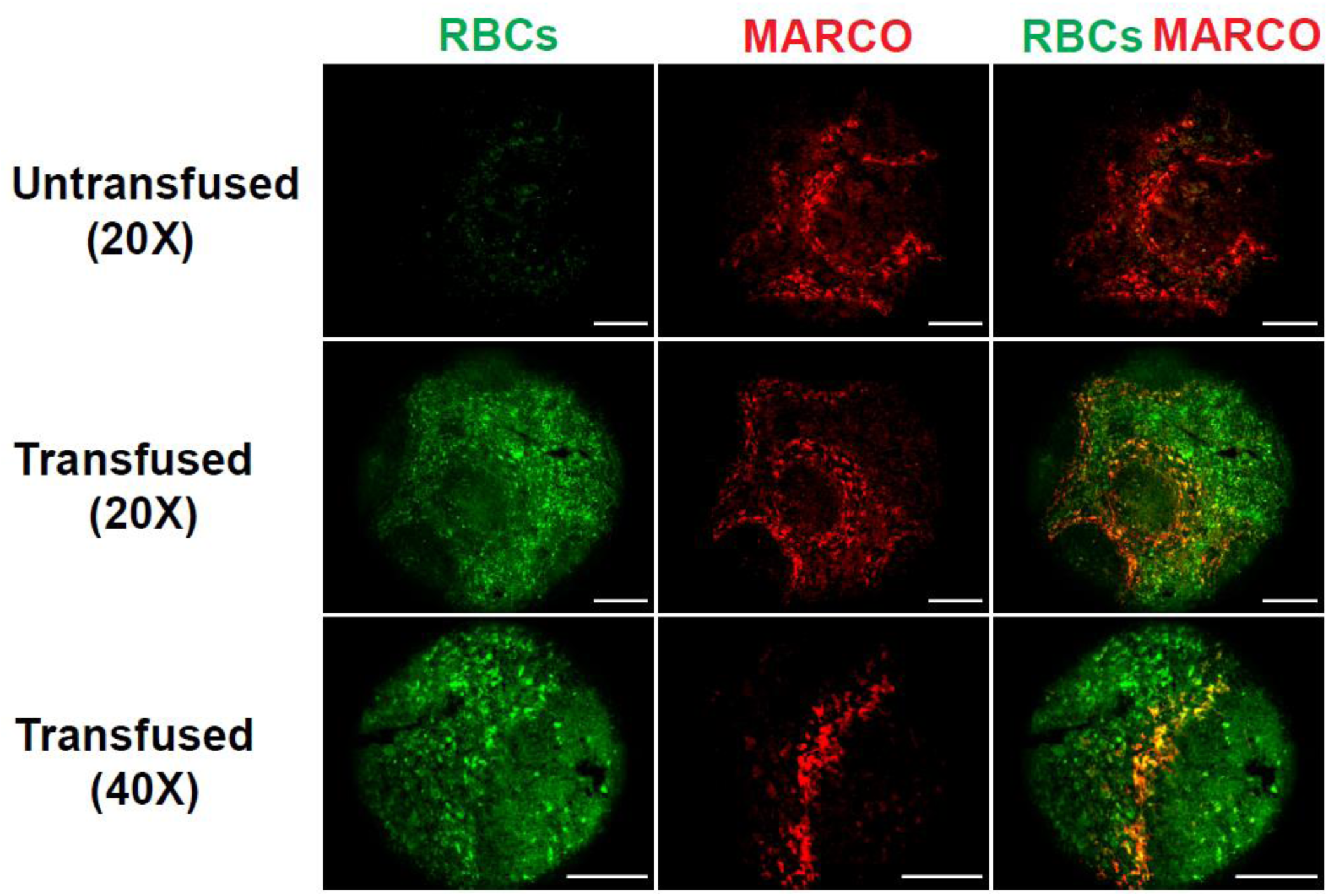
Stored RBCs localize to the splenic marginal zone macrophages. WT C57BL/6 mice, 8-12 weeks of age, were transfused with stored GFP expressing RBCs. Spleens were collected at 60min post-transfusion and processed for immunofluorescence microscopy. Splenic sections from stored GFP RBCs transfused mice were stained for MARCO^+^ MZMs (red). GFP RBCs were visualized directly (green). Splenic sections were imaged at the indicated magnifications (scale bar is 250μm).

**Figure 3.**
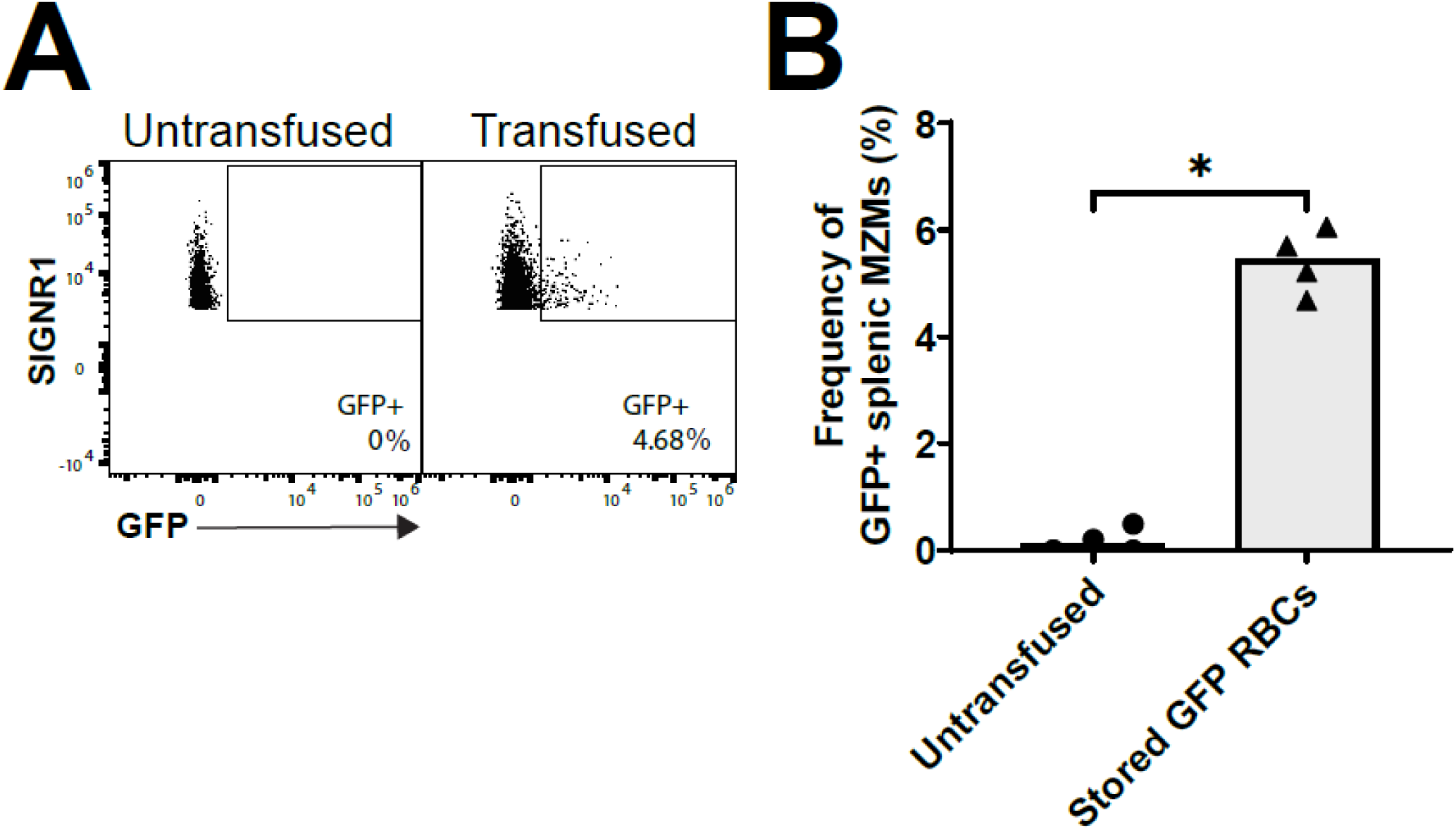
Stored RBCs are phagocytosed by splenic marginal zone macrophages post-transfusion. (A) Representative flow cytometry figures depicting the degree of phagocytosis of GFP RBCs (% GFP^+^) in SIGNR1^+^ splenic MZMs from mice that were transfused stored GFP RBCs or were untransfused. Extent of GFP^+^ RBC phagocytosis was measured at 60min post-transfusion. (B) Quantification of the frequency of GFP^+^ splenic MZMs from untransfused mice or mice transfused with stored RBCs at 60min post-transfusion. Data are representative of 3 independent experiments. P values were calculated using Kruskal-Wallis test followed by Mann-Whitney U test for post-hoc comparisons. **p** < 0.05, **\*\*p** < 0.01, **\*\*\*p** < 0.001and **ns p** > 0.05.

### LXRα-KO mice lack MZMs

Mice deficient in the transcription factor Liver X Receptor alpha (LXRα) have previously been reported to lack both MZMs as well as marginal metallophillic macrophages (MMMs)^23^. To confirm the reported macrophage phenotype in the LXRα-KO mice, we first stained splenic tissue sections from WT and LXRα-KO mice with the markers MARCO (MZMs) and MOMA (MMMs). We observed an absence of MARCO^+^ cellular staining in the marginal zone regions of splenic sections from LXRα-KO mice relative to WT mice, consistent with the published report (Figure 4A). Quantification of fluorescence intensity of MARCO staining showed significantly lower mean fluorescence intensity in LXRα-KO splenic sections compared to WT (Figure 4B). We did however continue to detect weak MARCO staining in conduit structures localized within the follicle that have previously been shown to be associated with FDC markers and to lie outside of the marginal zone^26^. We were surprised to observe MOMA^+^ staining in the marginal zone consistent with persistence of MMMs in our LXRα-KO mice spleens (Supplemental Figure 4A). This was unexpected given the previous published report^23^. This likely reflects a higher sensitivity of our fluorescent staining approach that is able to detect fewer MMM cells. To further validate the loss of MZMs, we used a recently described flow cytometric approach to quantify MZM levels that does not rely on MARCO staining^24^. In this approach, digestion of the spleen with an enzyme cocktail led to the identification of CD11b^+^F4/80^lo^Tim4^hi^ cells as splenic MZMs based on their high gene expression levels of MARCO and SIGNR1. Spleens from LXRα-KO mice showed a relative absence of CD11b^+^F4/80^lo^Tim4^hi^ cell population compared to WT spleens (Figure 4C, supplemental figure 4). Taken together, our data suggest that LXRα-KO mice specifically lack MZMs, while still retaining MMMs in the spleen.

**Figure 4.**
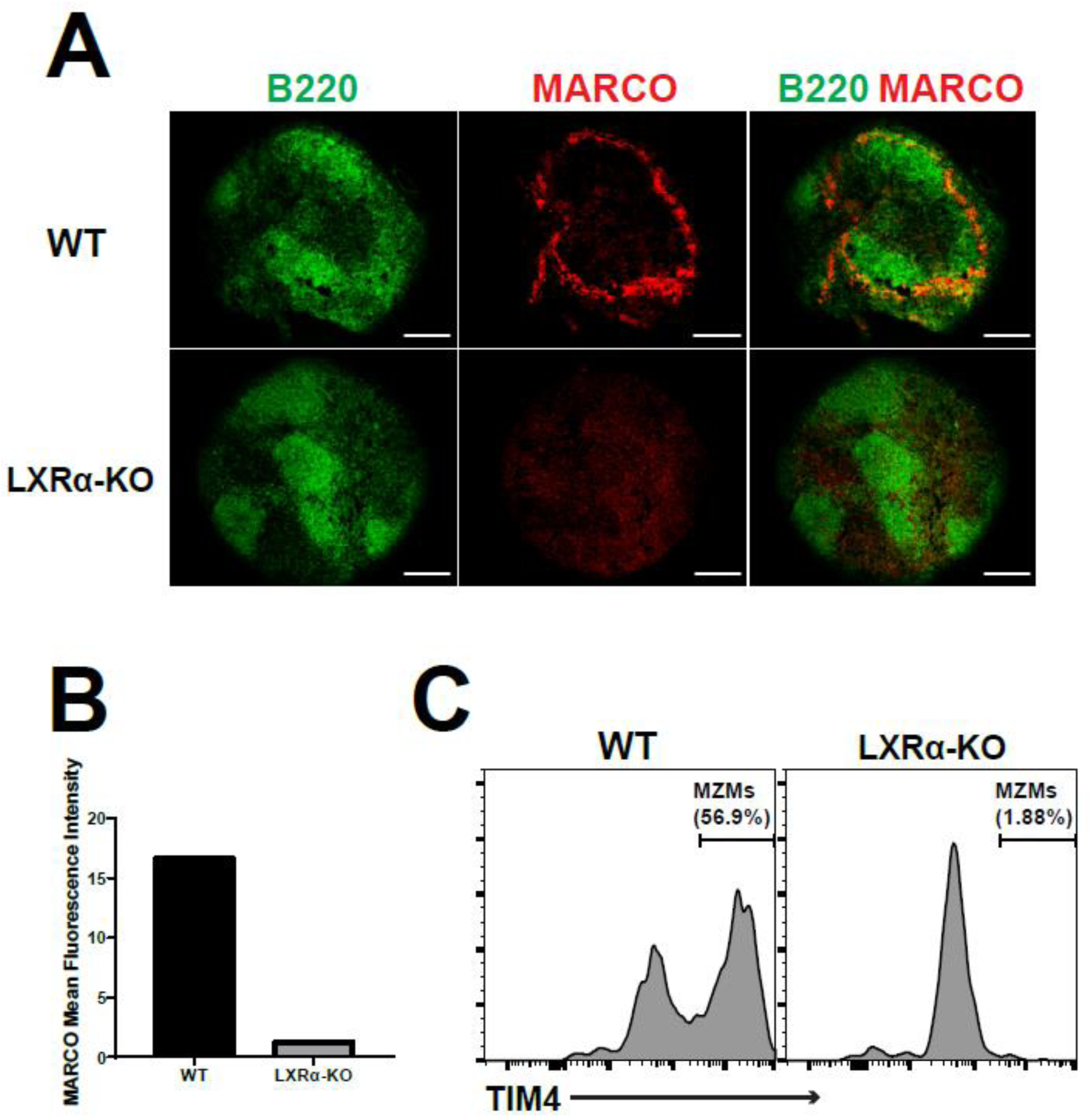
LXRa KO mice lack splenic MZMs. (A) Immunofluorescence images of splenic sections from WT and LXRα-KO mice, 8-12 weeks of age, stained with anti-MARCO antibody (red). Anti-B220 (green) was included in each image for visualization of B cell follicles. Images were taken at 20X magnification (Scale bar is 250μm). (B) Quantification of mean fluorescence intensity from MARCO staining for the images in (A). (C) Flow cytometry based identification of splenic MZMs in WT and LXRα-KO mice. Histograms show the frequency of MZMs (Tim4^hi^) in CD11b^+^F4/80^lo^ splenocytes from WT and LXRα-KO mice.

### LXRα-KO mice show reduced production of rapid phase inflammatory cytokines MCP-1 and KC, but not IL-6, in response to transfusion with stored HOD RBCs

To investigate the role of splenic marginal zone macrophages in the rapid production of inflammatory cytokines in response to transfused stored HOD RBCs, WT and LXRα-KO mice were transfused with 12-day stored HOD RBCs and sera were collected and compared for serum cytokine levels 90min post-transfusion. In 4 independent experiments, we found that, compared to WT mice, LXRα-KO mice had reduced serum levels of MCP-1 and KC, but no changes in IL-6 (Figure 5A-C, supplemental figure 5). Despite clear trends observed in all 4 experiments, statistical significance could not be reached in one of the replicates for MCP-1 (supplemental figure 5). Given this inconsistency, we decided to analyze all the replicates together using generalized estimating equation (GEE) statistical analysis to account for variance introduced from each independent study (supplementary methods). Our GEE analysis determined that on average LXRα-KO mice have 46% lower serum levels of MCP-1, 76% lower KC levels, and 0.45% higher IL-6 levels compared to WT mice in response to stored HOD RBC transfusion (Figure 5D). As LXRα-KO mice lack splenic MZMs, our results suggest a direct role of MARCO^+^ splenic MZMs in the production of rapid phase inflammatory cytokines MCP-1 and KC, but not IL-6.

**Figure 5.**
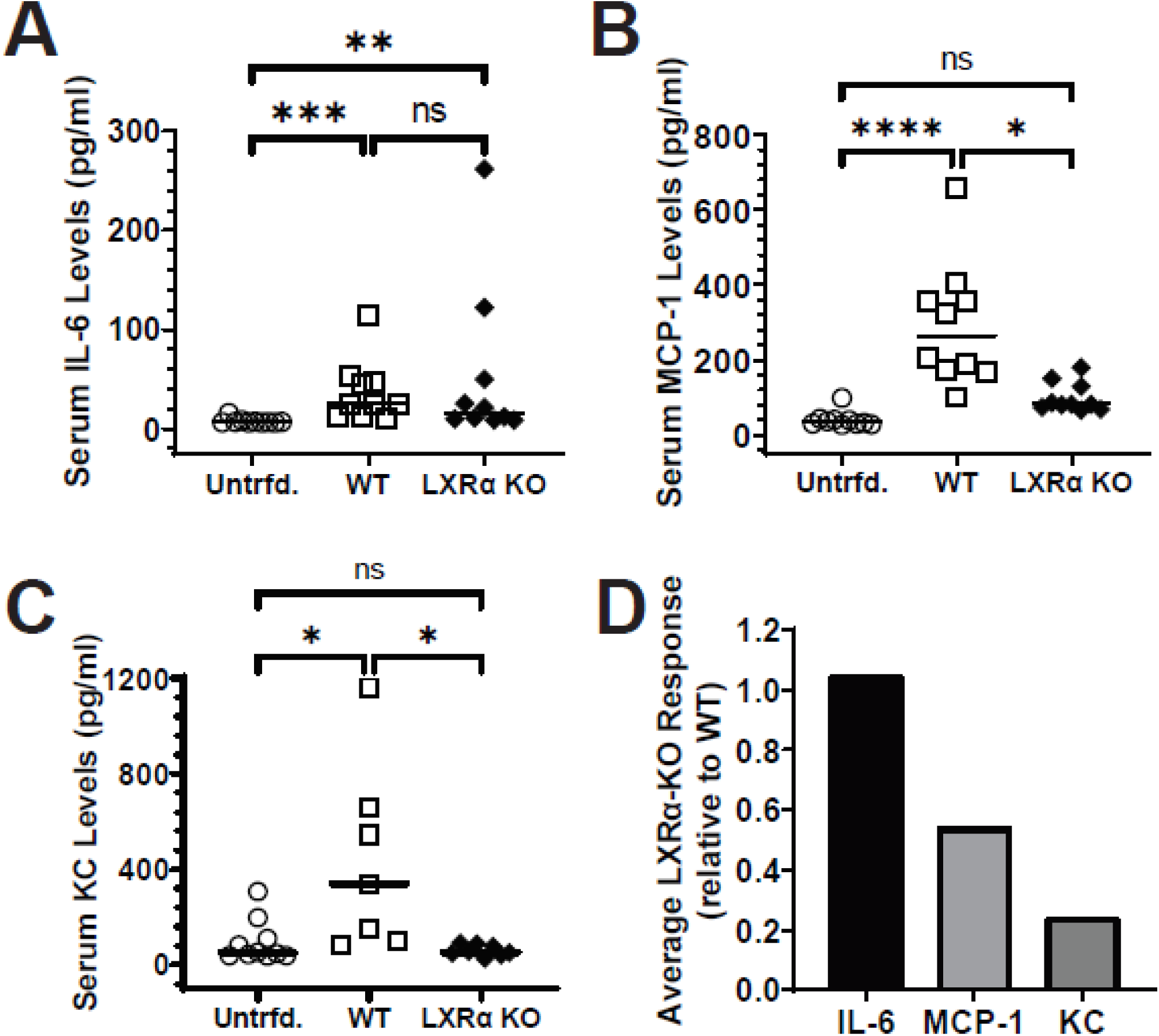
Absence of splenic marginal zone macrophages affects rapid production of inflammatory cytokines MCP-1 and KC, but not IL-6, in response to stored HOD RBC transfusion. WT and LXRα-KO naive recipients, 8-12 weeks of age, were transfused with 12-day stored HOD RBCs. Sera were collected 90min post-transfusion. Serum levels of MCP-1, KC, and IL-6 were measured and quantified through cytokine specific ELISAs including recombinant cytokine standard dilution curves. Serum levels of cytokines (A) IL-6, (B) KC, and (C) MCP-1 in WT and LXRα-KO mice. (D) Average cytokine response in LXRα-KO mice relative to WT mice as determined through GEE statistical modeling. Statistical significance was determined through non-parametric Kruskal-Wallis test followed by post-hoc comparisons using Mann-Whitney U test. **\*p** < 0.05, **\*\*p** < 0.01, **\*\*\*p** < 0.001and **ns p** > 0.05.

### MARCO^+^ splenic MZMs are not required for anti-HOD alloantibody generation in response to transfusion with stored HOD RBCs

To investigate the role of splenic MZMs in the regulation of adaptive immune responses against stored HOD RBCs, WT and LXRα-KO mice were transfused with 12-day stored HOD RBCs and sera were collected at days 7 and 14 post-transfusion to determine serum anti-HEL IgM and IgG levels respectively. We observed no significant difference between WT and LXRα-KO anti-HEL IgM levels 7-day post-transfusion with stored HOD RBCs as determined by endpoint ELISA titers (Figure 6A). Similarly, no statistically significant difference was observed between WT and LXRα-KO serum total anti-HOD IgG levels 14 days post-transfusion measured through flow cytometry cross-match (Figure 6B) or anti-HEL IgG levels determined through endpoint ELISA titers (Figure 6C). Despite the equivalent total anti-HEL IgG levels produced in WT and LXRα-KO mice, there could still be differences in the levels of individual IgG subtypes produced in the absence of splenic MZMs. We measured the serum levels of anti-HEL IgG1, IgG2b, IgG2c, and IgG3 antibodies 14 days post-transfusion with stored HOD RBCs. Similar to our results with anti-HEL IgM and total IgG antibody levels, no significant difference in the serum titers of anti-HEL IgG1, IgG2b, IgG2c, and IgG3 antibodies was observed as measured through endpoint ELISAs (Figure 7). These results show that the absence of splenic MZMs does not significantly affect RBC alloantibody production in response to stored HOD RBC transfusion.

**Figure 6.**
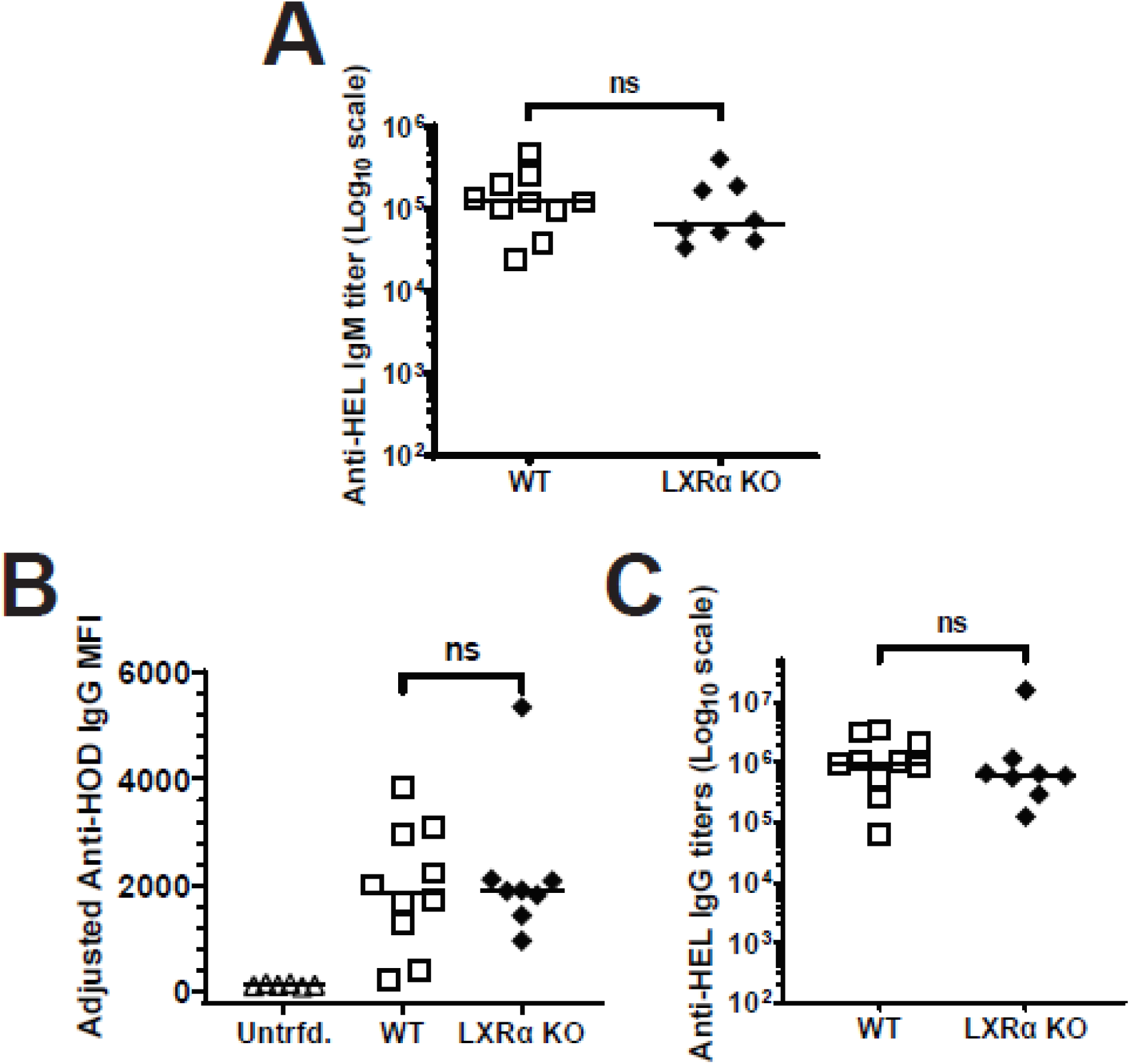
Splenic marginal zone macrophages are not required for production of anti-HOD alloantibody production in response to stored HOD RBC transfusion. WT and LXRα-KO mice, 8-12 weeks of age, were transfused with 12-day stored HOD RBCs and sera were collected at days 7 and 14 post-transfusion. Anti-HOD IgM serum titers were measured through anti-HEL ELISAs, and anti-HOD IgG alloantibody levels were determined using anti-HOD IgG RBC flow crossmatch and anti-HEL IgG ELISAs. (A) Serum anti-HEL IgM levels measured 7 days post-transfusion with stored HOD RBCs. (B) Flow crossmatch measuring serum anti-HOD IgG levels 14 days post-transfusion with stored HOD RBCs in WT and LXRα-KO mice depicted as adjusted MFI values, i.e., MFIs of recipient sera incubated with FVB RBCs subtracted from MFIs of sera incubated with HOD RBCs. (C) Serum anti-HEL total IgG titers measured 14 days post-transfusion with stored HOD RBCs through anti-HEL total IgG ELISA. Data are representative of 3-8 independent experiments. Statistical significance between experimental groups was determined through Kruskal-Wallis test and post-hoc comparisons were performed using Mann-Whitney U test. **\*p** < 0.05, **\*\*p** < 0.01, **\*\*\*p** < 0.001and **ns p** > 0.05.

**Figure 7.**
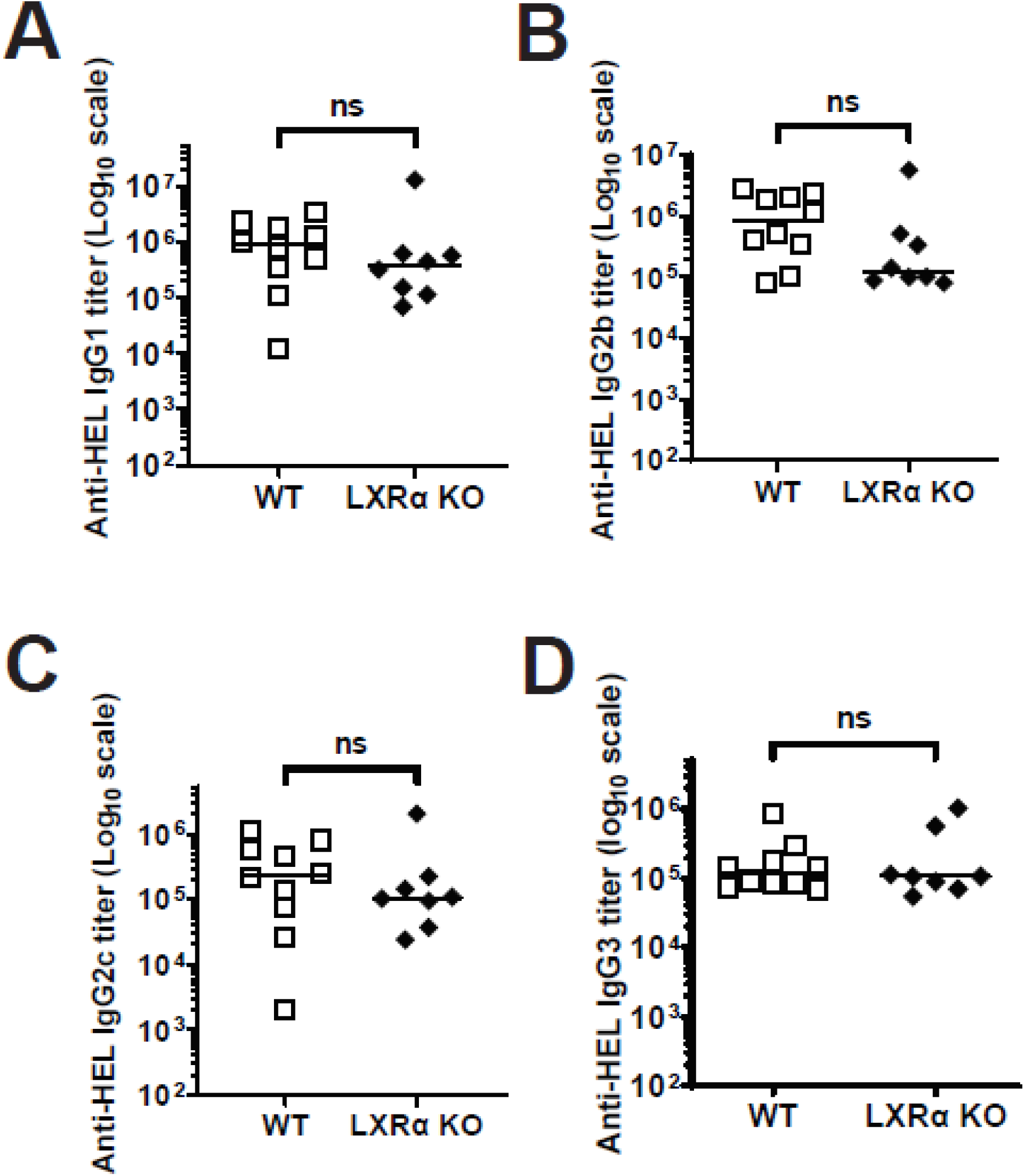
Absence of splenic MZMs does not affect the anti-HOD IgG subtype repertoire in response to transfused with stored HOD RBCs. WT and LXRα-KO mice, 8-12 weeks of age, were transfused with 12-day stored HOD RBCs and sera were collected 14 days post-transfusion. Anti-HOD IgG subtype serum titers were measured through anti-HEL IgG subtype specific ELISAs. Serum titers of anti-HEL IgG subtypes (A) IgG1, (B) IgG2b, (C) IgG2c, and (D) IgG3 in WT and LXRα-KO mice 14 days post-transfusion with stored HOD RBCs. Data are representative of 2-3 independent experiments. Non-parametric Kruskal-Wallis test followed by Mann-Whitney U tests for post-hoc comparisons was used to compare the experimental groups. **\*p** < 0.05, **\*\*p** < 0.01, **\*\*\*p** < 0.001and **ns p** > 0.05.

### Marginal Zone B cell dependent responses to a T-independent antigen are intact in LXRa KO mice

Given that MZMs are known to present antigen to MZBs, which are critical for RBC alloimmunization responses, we were surprised to find that a lack of MZMs did not affect alloantibody generation in LXRα-KO mice. Thus, we wanted to measure MZB cell function in LXRα-KO mice through immunization with TNP-Ficoll, a well characterized MZB cell dependent antigen. We compared the MZB cell frequencies and TNP-Ficoll responses in LXRα-KO mice to WT animals. LXRα-KO mice showed equivalent frequencies of MZB cells (defined as CD19^+^CD23^lo^CD21^hi^) compared to WT mice via flow cytometry (Fig. 8A-B, supplemental figure 6). Interestingly, no significant difference was found between titers of anti-TNP IgM and IgG between LXRα-KO and WT mice post-immunization with TNP-Ficoll (Fig. 8C-D). These results show that despite lacking splenic MZMs, LXRα-KO mice have an intact MZB cell compartment and normal responses to TNP-Ficoll.

**Figure 8.**
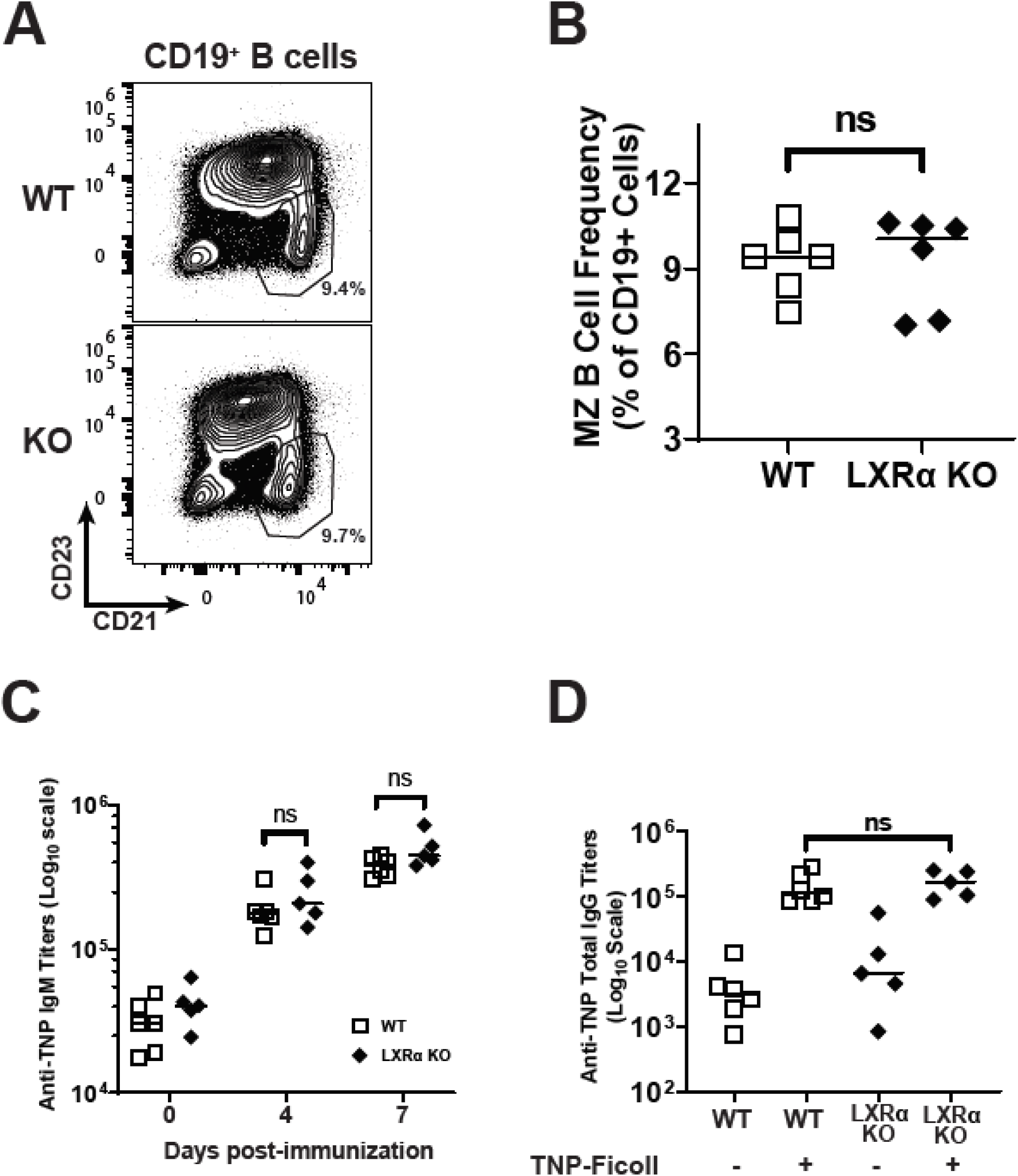
Lack of splenic MZMs in LXRα-KO mice does not affect MZ B cell frequencies or TNP-Ficoll responses. Comparison of splenic MZ B cell population levels between WT and LXRα-KO mice, 8-12 weeks of age, was performed through flow cytometry. (A) Representative flow cytometry contour plots showing frequency of MZ B cells (CD23^lo^CD21^hi^) within CD19^+^ B lymphocytes. (B) Quantification of MZ B cell frequencies in WT and LXRα-KO spleens. Responses of WT and LXRα-KO mice to TNP-Ficoll were determined through intravenous injection of 100μg of TNP-Ficoll in PBS per mouse, followed by sera collection and anti-TNP IgM and IgG responses determined at different time-points post-immunization. (C) Serum anti-TNP IgM titers in WT and LXRα-KO mice immunized with TNP-Ficoll at days 0, 4, and 7 post-immunization. (D) Serum anti-TNP total IgG levels in WT and LXRα-KO mice immunized with TNP-Ficoll at day 12 post-immunization. Anti-TNP titers were determined through anti-TNP endpoint titer ELISAs using TNP-BSA as the antigen. Data shown are representative to 2 independent experiments. Non-parametric Mann-Whitney U tests were used to determine statistical significance. **\*p** < 0.05, **\*\*p** < 0.01, **\*\*\*p** < 0.001and **ns p** > 0.05.

## DISCUSSION

All blood transfusions performed in patients utilize RBCs that have been stored for some period of time at 4^0^C. There is a great deal of variance in the storage times observed among transfused RBC units, with some units being stored less than 7 days while others are stored up to the FDA allowable 42 days prior to transfusion. Whether or not storage time impacts whether a given patient produces anti-RBC alloantibodies appears to depend on the clinical situation, with the length of RBC storage prior to transfusion failing to correlate with increased alloimmunization rates in the general population yet correlating with alloimmunization in patients with sickle cell disease^27–29^. In the HOD mouse model of anti-RBC alloimmunization, we and others have shown that storage of HOD RBCs at the extremes of FDA allowable storage results in significantly increased anti-RBC alloantibody production compared to freshly isolated RBCs^3,5,6,22^. Thus extended refrigerator storage of RBCs can alter their immunogenic properties in mice. By investigating the molecular and cellular mechanisms regulating immune response to stored RBCs, we hope to elucidate the key immunological pathways regulating anti-RBC alloimmunization.

Previous work has shown splenic tissue resident macrophages are involved in the rapid phase inflammatory cytokine production in response to stored RBCs^5^. In this report, we investigated the functional role of a specifically localized subpopulation of splenic macrophages in the alloimmunization responses to transfused stored RBCs, namely MARCO^+^ MZMs. We first showed that refrigerator stored RBCs co-localize with and are phagocytosed by splenic MZMs. Using LXRα-KO mice, we investigated immune responses to transfusion of stored HOD RBCs in the absence of splenic MZMs. We show that deletion of LXRα, which results in the loss of splenic MZMs, leads to a substantially lower production of rapid phase inflammatory cytokines MCP-1 and KC in response to transfused stored HOD RBCs. However, the absence of splenic MZMs does not affect the subsequent anti-HOD alloantibody generation.

Our clodronate results confirmed the requirement of phagocytes in general in the rapid removal of stored RBCs from circulation and generation of multiple inflammatory cytokines. Additionally, it demonstrates that that depletion of all phagocytes results in significantly lower levels of anti-HOD alloantibodies in response to stored HOD RBC transfusion. Thus, phagocytes are required for both innate and adaptive immune responses to stored RBCs, though it is certainly possible that different phagocyte populations might impact different stages of the immune response. Additionally, similar to the results in Larsson et al., refrigerator stored RBCs co-localize and are phagocytosed by MZMs. It is important to note that unlike ionomycin treated RBCs, not all stored RBCs co-localized with splenic MZMs. This is likely due to only a subset of stored RBCs upregulating PS and possibly other surface molecules, allowing them to be trapped and subsequently phagocytosed by splenic MZMs. Thus, refrigerator storage leads transfused RBCs to interact with the splenic marginal zone macrophage populations.

LXRα-KO mice produce substantially lower levels of MCP-1 and KC in response to stored HOD RBCs while IL-6 production is similar to WT mice. Several factors can explain the intact enhancement of alloantibody generation by stored HOD RBCs despite the decrease in rapid inflammatory responses. MCP-1 and KC production, albeit reduced, is still present in LXRα-KO mice. It is possible that despite reduced systemic levels of MCP-1 and KC, the local concentrations these cytokines in splenic tissue were sufficient to drive the enhancement of alloimmunization in response to stored HOD RBCs. We have previously shown IL-6 to be required for enhanced alloimmunization responses to stored HOD RBCs^22^, rapid production of which remains unaffected in LXRα-KO mice. As similar interrogations of the role of MCP-1 and KC remain to be performed, it is conceivable these cytokines may not be directly involved in regulating alloimmunization responses to stored HOD RBCs.

Characterization of LXRα-KO mice showed that they specifically lack splenic MZMs. An important function of splenic MZMs is to capture blood-borne antigens and present them to MZB cells, which were shown to be critical for alloimmunization responses to stored RBCs^9,15^. However, the absence of splenic MZMs in LXRα-KO mice did not significantly affect alloantibody generation in response to stored HOD RBCs. TNP-Ficoll responses also remained unaffected in the absence of splenic MZMs. Our results suggest that despite being able to capture and phagocytose stored RBCs, splenic MZMs are not necessary for alloantibody generation in response to stored HOD RBCs. We found that, unlike the previous report^23^, our LXRα-KO mice retained MMMs. MMMs are macrophages that line the opposite side of the marginal sinus closest to the T cell zone, and express the sialo-adhesion molecule CD169 recognized by the MOMA antibody^10,30^. MMMs are also able to sample both pathogens and apoptotic cells from blood in a similar manner to MZMs, but are not directly co-localized with MZBs. Instead, they are known to facilitate antigen trafficking to CD8^+^ dendritic cells in the white pulp^31^. Similar to MZMs, transfusion with stored RBC led to their co-localization with splenic MMMs. It is possible that in the absence of splenic MZMs, MMMs are able to present antigens to MZB cells suggesting a potential redundancy or compensation in antigen presentation to and activation of MZB cells. Stored RBCs were also shown to co-localize with MZB cells^9^, making it possible that MZB cells can get activated directly by stored RBCs without the need of antigen presenting cells. Further studies involving independent depletion of MMMs and combined depletion of both MMMs and MZMs are needed to fully characterize the role of these macrophages in the regulation of RBC alloimmunization.

The current study investigated the independent role of splenic MZMs in the enhancement of alloimmunization responses to stored RBCs. Although our results suggest a direct role for splenic MZMs in the production of inflammatory cytokines in response to stored RBCs, they were dispensable for alloantibody generation. Given the increased interaction of stored RBCs with phagocytic populations in the spleen, further studies delineating the independent roles of these cell populations in the regulation of immune response to stored RBCs would be of great interest and importance. The routine clinical use of refrigerator stored RBCs, and the relevance of storage length to alloimmunization rates in sickle cell patients, emphasizes the need for continued investigations of immune responses to transfused stored RBCs.

## ACKNOWLEDGEMENTS

The authors would like to thank all members of the Luckey lab for helpful discussions. The authors would also like to thank UVA Pathology microscopy core for their help with immunofluorescence imaging.

## SUPPLEMENTARY INFORMATION

### Supplementary Methods

#### Murine blood collection, processing, and storage

Blood from FVB.HOD or B6.GFP mice was collected directly into anticoagulant solution (CPDA-1, Boston Bioproducts) via cardiac puncture. Anticoagulant concentration was kept at 20% by volume. Blood was then leukoreduced using a neonatal leukoreduction filter (Acrodisc WBC syringe filter, Pall Corporation), centrifuged at 1200 × g for 10 minutes, and adjusted to a final hematocrit level of 75% (HOD RBCs) or 25% (B6.GFP RBCs) through removal of appropriate supernatant volume. Blood denoted as fresh was used for transfusions within 2 hours of processing. Stored blood was kept at 4°C for 12 days for FVB.HOD RBCs, and 14 days for B6.GFP RBCs.

#### Immunofluorescence microscopy

##### Immunofluorescence staining and imaging of splenic macrophage populations and splenic localization of transfused GFP RBCs

Spleens from WT and LXRα-KO mice were removed and fixed in 4% paraformaldehyde (Electron Microscopy Sciences, Hatfield, PA) for 2 hours, suspended in 30% sucrose overnight, embedded in optimal cutting temperature (OCT) compound, and snap-frozen with 2-methylbutane (Sigma-Aldrich, St. Louis, MO) and liquid nitrogen. Splenic sections, 10μm thick, were blocked with 10% normal goat serum (NGS) in PBS and then stained with primary antibodies overnight at 4°C and secondary antibodies at RT for 1h in 5% NGS in PBS. Primary antibodies included purified anti-mouse CD169 (MOMA, 1:100, Bio-Rad) and purified anti-mouse MARCO (1:50, LifeSpan Biosciences). Goat anti-rabbit IgG or goat anti-rat IgG conjugated to AlexaFluor 555 were used as secondary antibodies (1:200, Invitrogen).

For GFP RBC splenic localization studies, spleens from WT mice transfused with stored B6.GFP RBCs were removed 60min post-transfusion and processed for imaging as mentioned above. Spleens from transfused mice were stained for MARCO and MOMA. GFP RBCs were visualized directly.

Splenic sections were visualized on an Olympus BX40 fluorescence microscope (Olympus Life Science) and images were processed and quantified using ImageJ software.

#### Flow cytometry

##### Erythrocyte phagocytosis assay

To determine phagocytosis of RBCs by splenic macrophage populations, spleens from WT C57BL/6 mice transfused with stored B6.GFP RBCs were collected 60min post-transfusion. Spleens were processed into single cell suspensions, followed by lysis of non-phagocytosed RBCs. Splenocytes were then stained for MZMs (SIGNR1) in FACS buffer (PBS + 2%FBS + 0.5% BSA + 0.1% NaN_3_). Frequency of GFP^+^ cells were determined through flow cytometry. Reagents for staining included, anti-SIGNR1 APC and fixable live dead dye efluor-780 from ThermoFisher Scientific, and anti-CD11b PE-Cy7, anti-CD3 APC-Cy7, anti-B220 APC-Cy7, and anti-TER119 PerCP-Cy5.5 from Biolegend.

##### MZ B cell staining

Spleens from WT and LXRα-KO mice were processed into single cell suspensions. Splenocytes were then stained with anti-mouse CD19 BV421, CD23 PE, CD21 PE-Cy7, CD3 FITC, and 7-AAD for dead cell exclusion in FACS buffer on ice for 20min. All staining reagents were purchased from Biolegend.

##### MZM staining

Spleens were processed for MZM staining as described previously^24^. Splenocytes were first stained with efluor-780 fixable viability dye (ThermoFisher Scientific) followed by staining with anti-CD3, anti-CD19, anti-CDNK1.1, anti-Ly6G, anti-SiglecF all conjugated to FITC (Biolegend), anti-F4/80 conjugated to brilliant violet 421 (Biolegend), and anti-CD11b conjugated to PE-Cy7 (Biolegend).

For all flow cytometry experiments, samples were acquired using the Attune NXT flow cytometer (ThermoFisher Scientific) and data were analyzed using Flowjo (BD Biosciences) and GraphPad Prism.

#### Serum cytokine and antibody measurements

Sera from recipient mice were collected via submandibular bleeding 90 minutes post-transfusion for cytokine measurements. Serum MCP-1, KC, and IL-6 levels were measured using cytokine specific ELISA kits following the manufacturer’s instructions. Mouse MCP-1 and IL-6 ELISA kits were purchased from Invitrogen (ThermoFisher Scientific, Waltham MA), and mouse KC ELISA kit was purchased from R&D Systems (BioTechne, Minneapolis, MN).

#### Serum Anti-HOD alloantibody flow cytometry crossmatch

FVB.HOD and FVB/NJ RBCs were collected in CPDA-1, washed in FACS buffer and incubated with recipient sera collected 14 days post-transfusion at a 1:8 dilution for 30min at room temperature. RBCs were then stained with phycoerythrin-conjugated goat anti-mouse IgG secondary antibody (ThermoFisher Scientific, St. Louis, MO) for 30min at room temperature. Following secondary staining, RBCs were washed, resuspended in FACS buffer, and data were collected through flow cytometry on the Attune NxT instrument (ThermoFisher Scientific, St. Louis, MO). Flow cytometry data were analyzed using FlowJo (BD Biosciences) and MS Excel. For each individual recipient mouse, median fluorescence intensities (MFIs) measured from FVB/NJ RBCs were subtracted from MFIs from FVB.HOD RBCs to obtain anti-HOD specific median fluorescence signals, referred to as adjusted MFIs. Data plotting, figure construction, and statistical analyses were performed on GraphPad Prism.

#### Serum Anti-HEL endpoint titer ELISAs

High protein-binding ELISA plates (Corning 9018) were coated overnight at 4°C with 10µg/ml Hen Egg Lysozyme protein (Sigma) in PBS. Plates were then washed (0.05% Tween in PBS) and incubated with blocking buffer (2% BSA and 0.05% Tween-20 in PBS) for 1h at 37°C or overnight at 4°C. Sera from recipient mice were serially diluted (starting at 1:50) in block buffer by a factor of 4 each time for 12 total dilutions and were incubated on the coated plates for 2hr at RT. Plates were washed and incubated with a horseradish peroxidase (HRP)-conjugated anti-mouse antibody isotype specific secondary antibody at 1:5000 dilution for 1 hour at RT. Secondary antibodies included HRP-conjugated goat anti-mouse IgM, IgG, IgG1, IgG2b, IgG2c, and IgG3 (Jackson Immunoresearch). Wells were then developed using TMB substrate (SeraCare) for 10 minutes and quenched with 2N H_2_SO_4_. Optical densities were observed at 450nm on SpectraMax Plus microplate spectrophotometer and end-point titer values were calculated using GraphPad Prism. Values were calculated by interpolation of the cutoff point where optical density values for each sample were plotted against serum dilution using a “plateau followed by one phase decay” model. Cutoff values were defined as the signal from blank background wells plus three standard deviations.

#### Stored HOD RBC post-transfusion recovery analysis

Blood, ∼5-10ul, from recipient mice transfused with 12-day stored HOD RBCs was collected via tail snip 24h post-transfusion directly into 100μl CPDA-1 and washed with FACS buffer. The RBCs were then stained with mouse anti-human duffy blood group antibody (MIMA29, New York Blood Center) at a 1:100 dilution in FACS buffer and incubated at room temperature for 30min. RBCs were then washed and stained with PE-conjugated goat anti-mouse IgG secondary antibody (ThermoFisher Scientific) at a 1:200 dilution in FACS buffer for 30min at room temperature. RBCs were washed for a final time, resuspended in FACS buffer and acquired using the Attune NXT flow cytometer (ThermoFisher Scientific) for the presence and quantification of circulating HOD+ RBCs in recipient mice. Data were analyzed using flowJO (BD Biosciences), and GraphPad Prism.

#### T independent antigen immunization and serum immunoglobulin measurements

WT C57BL/6 and LXRα-KO were immunized with 100μg of hapten 2,4,6-Trinitrophenol conjugated Ficoll (TNP-Ficoll, Biosearch Technologies) in PBS through intravenous injections via the retro-orbital sinus, and sera were collected at days 4, 7, and 12 post-immunization. Anti-TNP IgM, and total IgG were determined through anti-TNP ELISAs using TNP-BSA (Biosearch Technologies) as the antigen and following the same protocol and reagents as described above for anti-HEL endpoint titer ELISAs.

#### Generalized estimation equation analysis for comparison of cytokine production post-transfusion

GEE analysis was performed on combined cytokine production data from multiple replicates to determine the effect of LXRα deletion while accounting for technical and biological variability resulting from independent experiments. Given the log-normal distribution of cytokine responses, a Gaussian model with a log-link function, exchangeable correlation and robust standard errors was fitted to the data. The effects of LXRα deletion on cytokine production was determined from the GEE model fit and interpreted as an average fold-change compared to cytokine responses of WT mice. All analyses were performed using R.

## Supplementary figure legends

**Supplementary figure 1.**
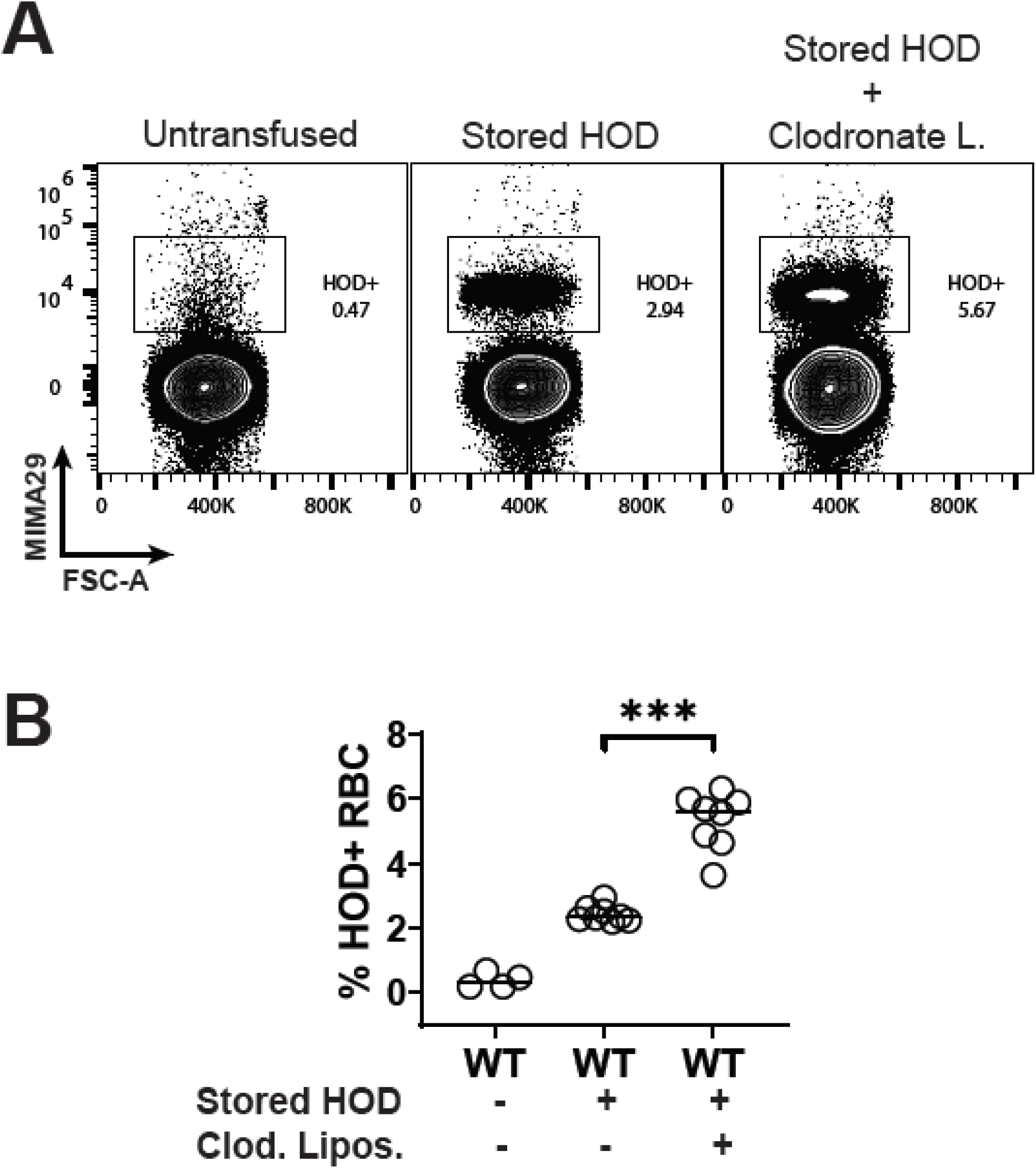
Increased post-transfusion survival of stored HOD RBCs in phagocyte depleted mice. WT C57BL/6 mice were treated with liposomal clodronate 48-72h prior to transfusion with stored HOD RBCs. Frequency of HOD^+^ RBCs in circulation post-transfusion was measured via flow cytometry. (A) Representative flow cytometry contour plots depicting staining of HOD^+^ RBCs in circulation 24h post-transfusion in mice treated with clodronate liposomes or left untreated. (B) Frequency of transfused HOD^+^ RBCs, as a percentage of total RBCs, in recipient mice 24h post-transfusion with stored HOD RBCs. Data shown are representative of 3 independent experiments. **\*p** < 0.05, **\*\*p** < 0.01, **\*\*\*p** < 0.001 and **ns p** > 0.05.

**Supplementary figure 2.**
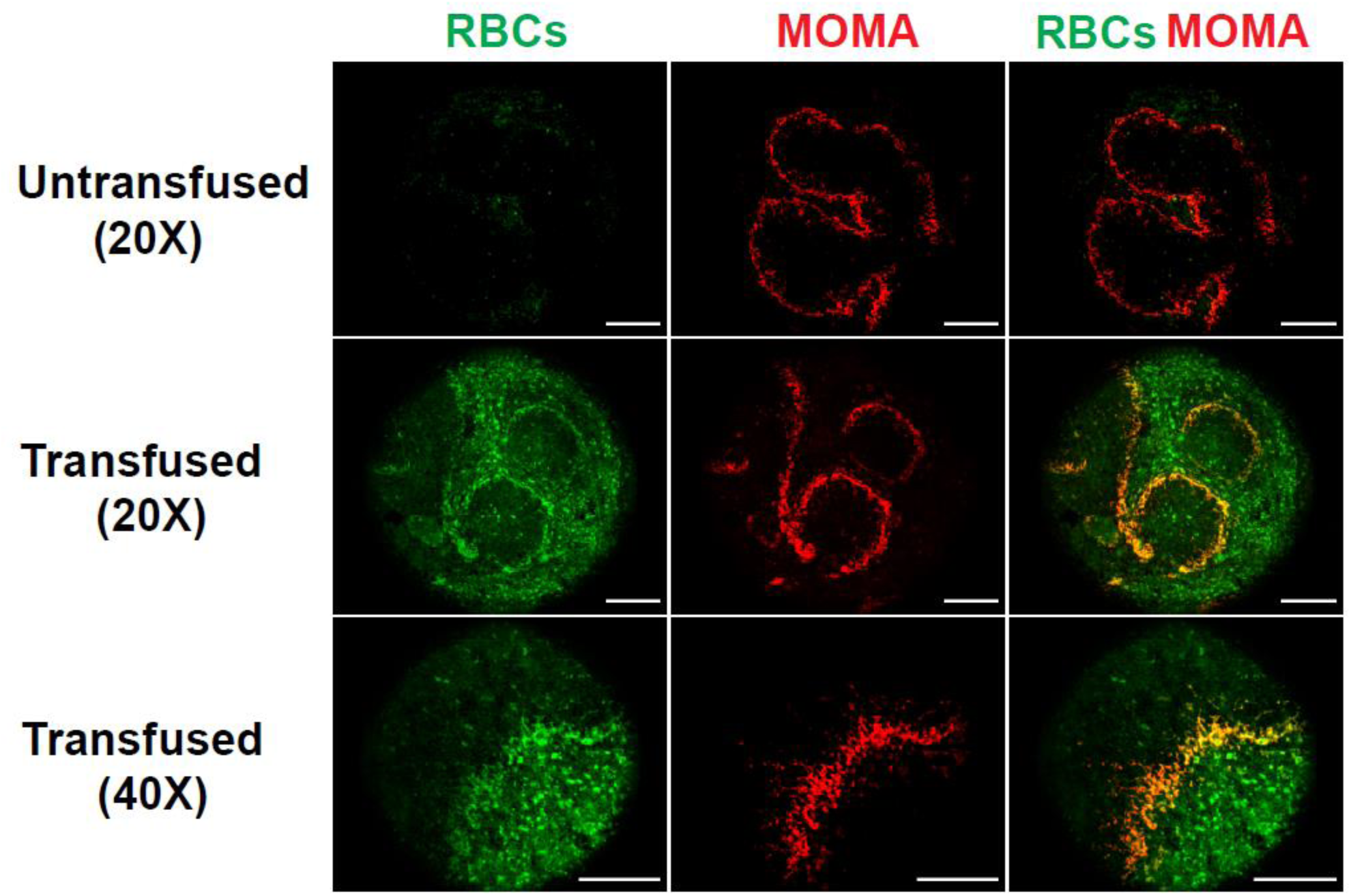
Stored RBCs localize to marginal metallophilic macrophages. WT C57BL/6 mice, 8-12 weeks of age, were transfused with stored GFP^+^ RBCs. Spleens were collected at 60min post-transfusion and imaged via immunofluorescence microscopy. Splenic sections from stored GFP RBCs transfused mice were stained for MOMA^+^ MMMs (red). GFP RBCs were visualized directly (green). Splenic sections were imaged at the indicated magnifications (scale bar is 250μm).

**Supplementary figure 3.**
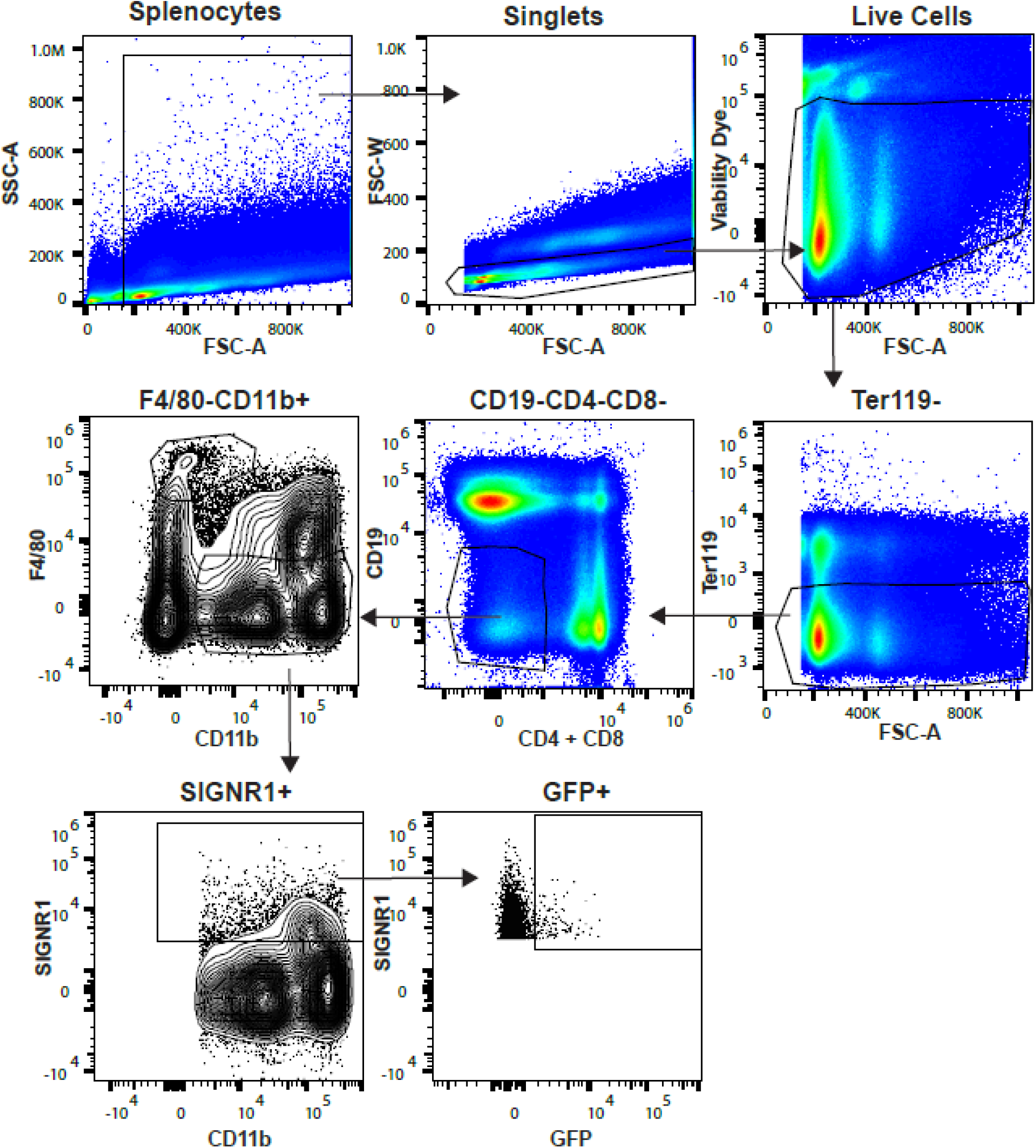
Flow cytometry gating strategy for determination of stored GFP RBCs phagocytosis by splenic MZMs post-transfusion. Samples were first gated on all splenocytes, followed by elimination of doublets, dead cells, and extracellular RBCs. The cells were then gated on non-lymphocytes, followed by F4/80^-^ macrophages (to exclude red pulp macrophages), and finally gating on SIGNR1^+^ splenic MZMs.

**Supplementary figure 4.**
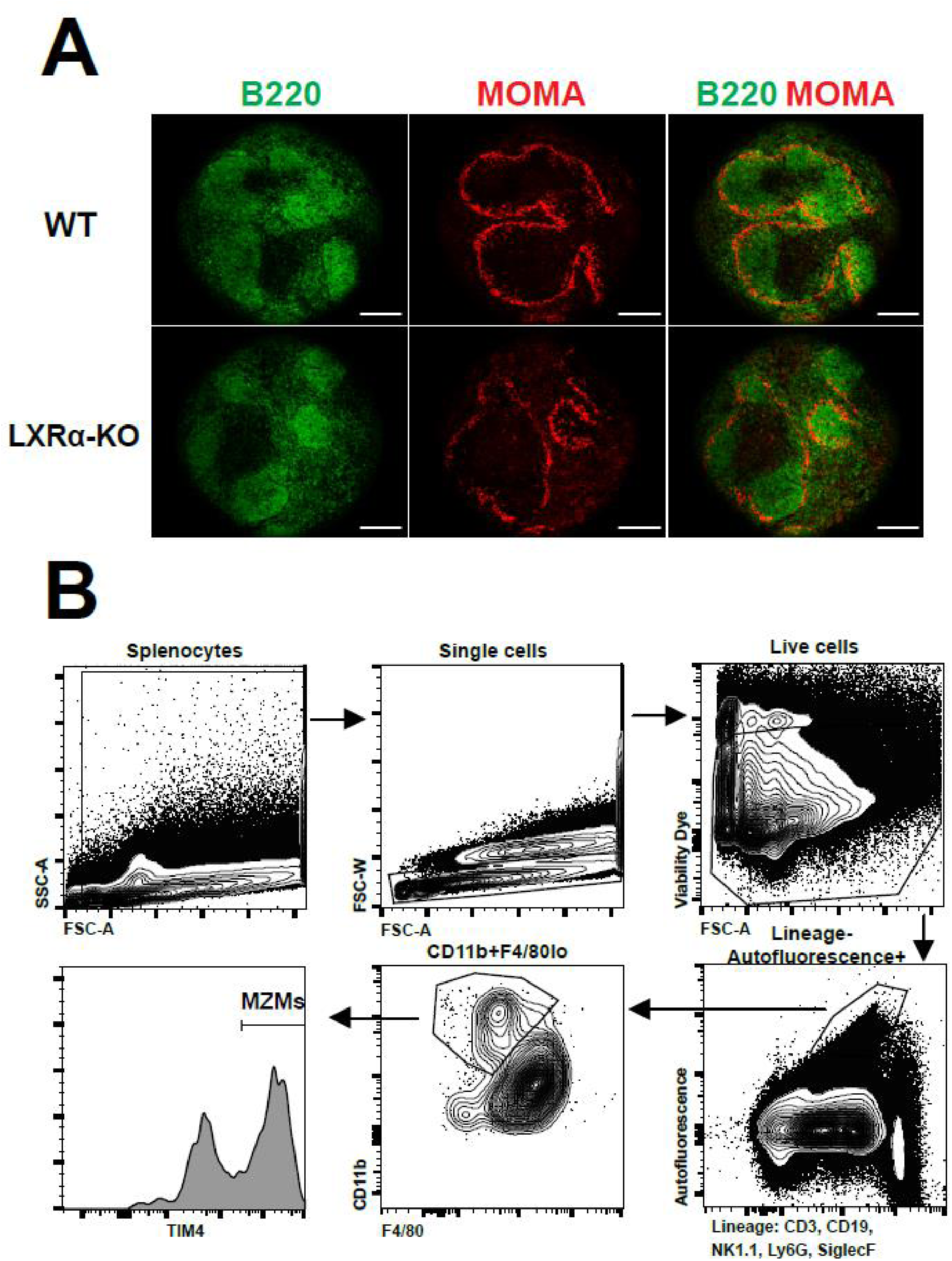
LXRα-KO mice retain splenic MMMs, and gating strategy for flow cytometry based identification of splenic MZMs. (A) Immunofluorescence images of splenic sections from WT and LXRα-KO mice, 8-12 weeks of age, stained with anti-MOMA antibody (red) and anti-B220 (green) for visualization of B cell follicles. Images were taken at 20X magnification (Scale bar is 250μm). (B) After spleen processing and staining, samples were first gates on all splenocytes, followed by exclusion of doublets and dead cells. Live cells were gated on autofluorescencent cells that were negative for all lineage markers, followed by gating on CD11b^+^F4/80^lo^ cells. Splenic MZMs were finally identified as Tim4^hi^ cells within the CD11b^+^F4/80^lo^ gated cells.

**Supplementary figure 5.**
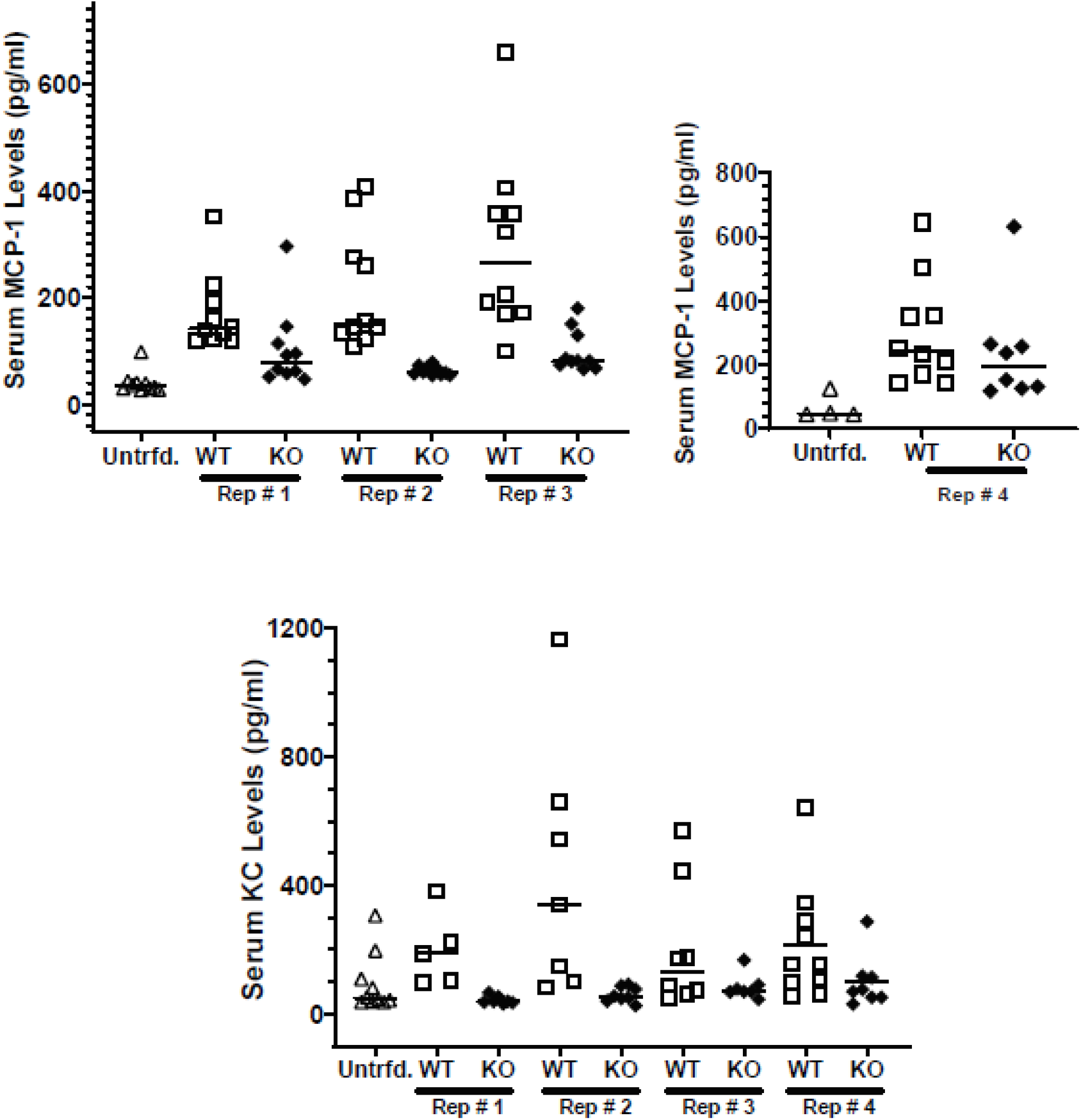
LXRα-KO mice show lower rapid production of MCP-1 and KC, but not IL-6, post-transfusion with stored HOD RBCs. WT C57BL/6 and LXRα-KO mice were transfused with stored HOD RBCs, and serum levels of IL-6, MCP, and KC were measured 90min post-transfusion. Serum cytokine levels were measured via cytokine specific ELISAs. Figures depicting serum cytokine levels in all 4 independent experiments for (A) IL-6, (B) MCP-1, and (C) KC.

**Supplementary figure 6.**
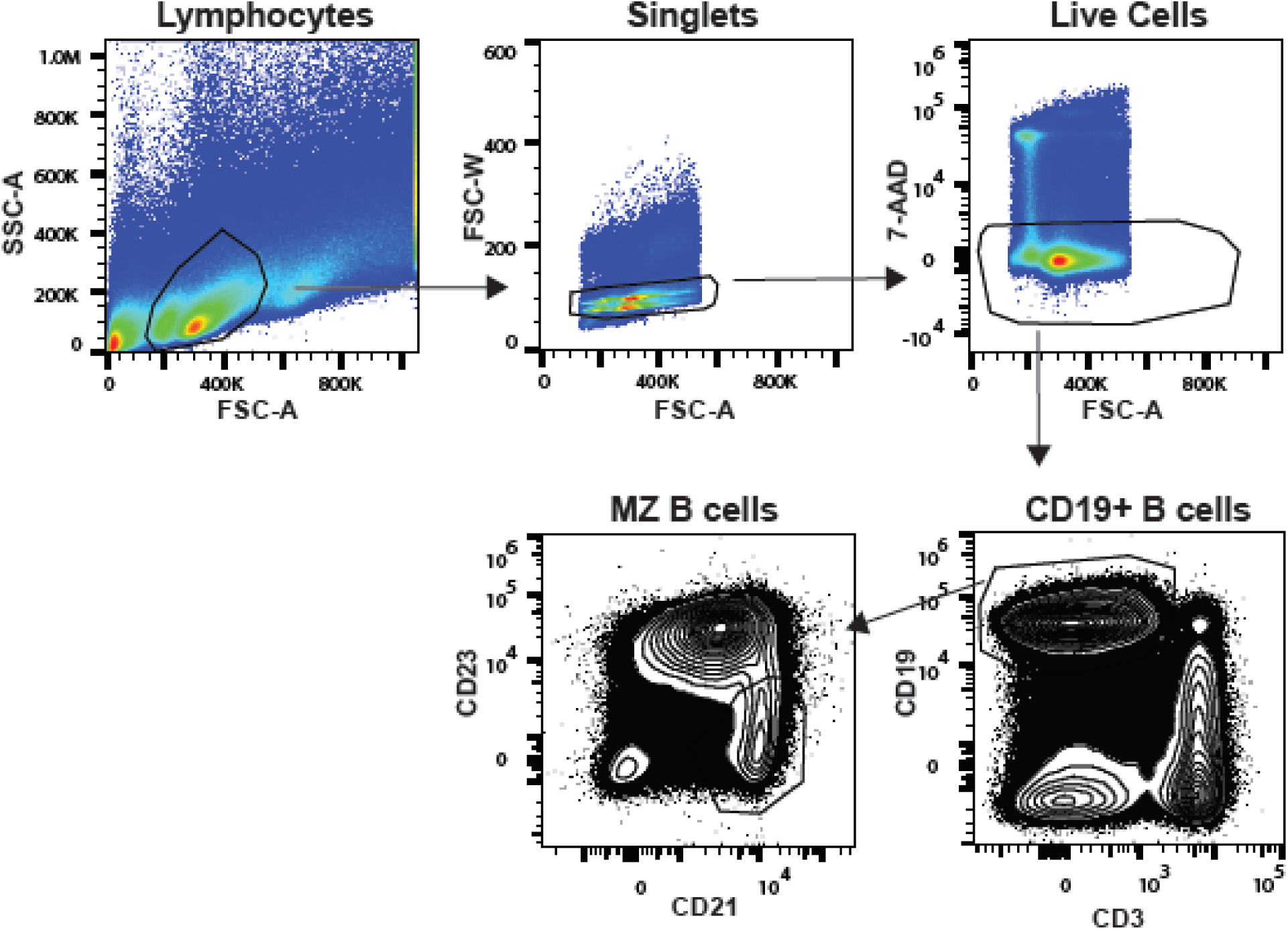
LXRα-KO mice have equivalent MZ B cell frequencies compared to WT mice. Flow cytometry gating strategy for determination of MZ B cell frequencies in the spleens of WT and LXRα-KO mice. Samples were first gated on lymphocytes followed by exclusion of doublets and dead cells. Live single cells were then gated on CD19^+^ B lymphocytes, followed by gating on CD21^hi^CD23^lo^ MZ B cells.

## Notes

### Competing Interest Statement

The authors have declared no competing interest.

### Summary of Updates

The funding sources have been added and no further edits were made.

### Funding

- NIH Common Fund: R01HL134691, P01HL169552

## REFERENCES

1. Tormey CA, Hendrickson JE. Transfusion-related red blood cell alloantibodies: induction and consequences. Blood. 2019 Apr 25;133(17):1821–30.

2. Zimring JC, Hudson KE. Cellular immune responses in red blood cell alloimmunization. Hematology American Society of Hematology Education Program. 2016 Dec 2;2016(1):452–6.

3. Hod EA, Zhang N, Sokol SA, Wojczyk BS, Francis RO, Ansaldi D, et al. Transfusion of red blood cells after prolonged storage produces harmful effects that are mediated by iron and inflammation. Blood. 2010 May 27;115(21):4284–92.

4. Arneja A, Salazar JE, Jiang W, Hendrickson JE, Zimring JC, Luckey CJ. Interleukin-6 receptor-alpha signaling drives anti-RBC alloantibody production and T-follicular helper cell differentiation in a murine model of red blood cell alloimmunization. Haematologica. 2016 Nov;101(11):e440–4.

5. Wojczyk BS, Kim N, Bandyopadhyay S, Francis RO, Zimring JC, Hod EA, et al. Macrophages clear refrigerator storage-damaged red blood cells and subsequently secrete cytokines in vivo, but not in vitro, in a murine model: Transfusion-Associated Inflammation. Transfusion. 2014 Dec;54(12):3186–97.

6. Hendrickson JE, Hod EA, Spitalnik SL, Hillyer CD, Zimring JC. IMMUNOHEMATOLOGY: Storage of murine red blood cells enhances alloantibody responses to an erythroid-specific model antigen: ALLOIMMUNOGENICITY OF STORED MURINE RBCs. Transfusion. 2009 Nov 9;50(3):642–8.

7. Hendrickson JE, Saakadze N, Cadwell CM, Upton JW, Mocarski ES, Hillyer CD, et al. The spleen plays a central role in primary humoral alloimmunization to transfused mHEL red blood cells. Transfusion. 2009 Aug;49(8):1678–84.

8. Evers D, van der Bom JG, Tijmensen J, de Haas M, Middelburg RA, de Vooght KMK, et al. Absence of the spleen and the occurrence of primary red cell alloimmunization in humans. Haematologica. 2017 Aug;102(8):e289–92.

9. Zerra PE, Patel SR, Jajosky RP, Arthur CM, McCoy JW, Allen JWL, et al. Marginal Zone B Cells Mediate a CD4 T Cell Dependent Extrafollicular Antibody Response Following RBC Transfusion in Mice. Blood. 2021 Apr 19;

10. Cyster JG. B cells on the front line. Nature Immunology. 2000 Jul;1(1):9–10.

11. Lewis SM, Williams A, Eisenbarth SC. Structure and function of the immune system in the spleen. Science Immunology. 2019 01;4(33).

12. Vanderkerken M, Maes B, Vandersarren L, Toussaint W, Deswarte K, Vanheerswynghels M, et al. TAO-kinase 3 governs the terminal differentiation of NOTCH2-dependent splenic conventional dendritic cells. Proceedings of the National Academy of Science U S A. 2020 Dec 8;117(49):31331–42.

13. You Y, Myers RC, Freeberg L, Foote J, Kearney JF, Justement LB, et al. Marginal Zone B Cells Regulate Antigen Capture by Marginal Zone Macrophages. Journal Immunology. 2011 Feb 15;186(4):2172–81.

14. Aichele P, Zinke J, Grode L, Schwendener RA, Kaufmann SHE, Seiler P. Macrophages of the Splenic Marginal Zone Are Essential for Trapping of Blood-Borne Particulate Antigen but Dispensable for Induction of Specific T Cell Responses. Journal Immunology. 2003 Aug 1;171(3):1148–55.

15. den Haan JMM, Kraal G. Innate Immune Functions of Macrophage Subpopulations in the Spleen. Journal of Innate Immunolgy. 2012;4(5–6):437–45.

16. Antonelou MH, Seghatchian J. Insights into red blood cell storage lesion: Toward a new appreciation. Transfusion and Apheresis Science. 2016 Dec;55(3):292–301.

17. Tissot J-D, Bardyn M, Sonego G, Abonnenc M, Prudent M. The storage lesions: From past to future. Transfusion Clinique et Biologique. 2017 Sep;24(3):277–84.

18. Zimring JC. Established and theoretical factors to consider in assessing the red cell storage lesion. Blood. 2015 Apr 2;125(14):2185–90.

19. Lang E, Pozdeev VI, Xu HC, Shinde PV, Behnke K, Hamdam JM, et al. Storage of Erythrocytes Induces Suicidal Erythrocyte Death. Cellular Physiology and Biochemistry. 2016;39(2):668–76.

20. Larsson A, Hult A, Nilsson A, Olsson M, Oldenborg P-A. Red blood cells with elevated cytoplasmic Ca ^2+^ are primarily taken up by splenic marginal zone macrophages and CD207+ dendritic cells: RBC CLEARANCE IN THE SPLENIC MARGINAL ZONE. Transfusion. 2016 Jul;56(7):1834–44.

21. Desmarets M, Cadwell CM, Peterson KR, Neades R, Zimring JC. Minor histocompatibility antigens on transfused leukoreduced units of red blood cells induce bone marrow transplant rejection in a mouse model. Blood. 2009 Sep 10;114(11):2315–22.

22. Arneja A, Salazar JE, Jiang W, Hendrickson JE, Zimring JC, Luckey CJ. Interleukin-6 receptor-alpha signaling drives anti-RBC alloantibody production and T-follicular helper cell differentiation in a murine model of red blood cell alloimmunization. Haematologica. 2016 Nov;101(11):e440–4.

23. A-Gonzalez N, Guillen JA, Gallardo G, Diaz M, de la Rosa JV, Hernandez IH, et al. The nuclear receptor LXRα controls the functional specialization of splenic macrophages. Nature Immunology. 2013 Aug;14(8):831–9.

24. Fujiyama S, Nakahashi-Oda C, Abe F, Wang Y, Sato K, Shibuya A. Identification and isolation of splenic tissue-resident macrophage sub-populations by flow cytometry. International Immunology. 2019 Feb 6;31(1):51–6.

25. van Rooijen N, Hendrikx E. Liposomes for specific depletion of macrophages from organs and tissues. Methods Molecular Biology. 2010;605:189–203.

26. Kellermayer Z, Fisi V, Mihalj M, Berta G, Kóbor J, Balogh P. Marginal Zone Macrophage Receptor MARCO Is Trapped in Conduits Formed by Follicular Dendritic Cells in the Spleen. Journal of Histochemistry & Cytochemistry. 2014 Jun;62(6):436–49.

27. Desai PC, Deal AM, Pfaff ER, Qaqish B, Hebden LM, Park YA, et al. Alloimmunization is associated with older age of transfused red blood cells in sickle cell disease. American Journal of Hematology. 2015 Aug;90(8):691–5.

28. Zalpuri S, Schonewille H, Middelburg R, van de Watering L, de Vooght K, Zimring J, et al. Effect of storage of red blood cells on alloimmunization. Transfusion. 2013 Nov;53(11):2795–800.

29. Dinardo CL, Fernandes FLA, Sampaio LR, Sabino EC, Mendrone A. Transfusion of older red blood cell units, cytokine burst and alloimmunization: a case-control study. Brazilian Journal of Hematology and Hemotherapy. 2015 Oct;37(5):320–3.

30. Lewis SM, Williams A, Eisenbarth SC. Structure and function of the immune system in the spleen. Science Immunology. 2019 01;4(33).

31. Backer R, Schwandt T, Greuter M, Oosting M, Jüngerkes F, Tüting T, et al. Effective collaboration between marginal metallophilic macrophages and CD8+ dendritic cells in the generation of cytotoxic T cells. Proceedings of the National Academy of Science U S A. 2010 Jan 5;107(1):216–21.

